# Collective strategy for obstacle navigation during cooperative transport by ants

**DOI:** 10.1101/061036

**Authors:** Helen F. McCreery, Zachary A. Dix, Michael D. Breed, Radhika Nagpal

## Abstract

Group cohesion and consensus have primarily been studied in the context of discrete decisions, but some group tasks require making serial decisions that build on one another. We examine such collective problem solving by studying obstacle navigation during cooperative transport in ants. In cooperative transport, a group of ants works together to move a large object back to their nest. We blocked cooperative transport groups of *Paratrechinal longicornis* with obstacles of varying complexity, analyzing groups trajectories to infer what kind of strategy the ants employed. Simple strategies require little information, but more challenging, robust strategies succeed with a wider range of obstacles. We found that transport groups use a stochastic strategy that leads to efficient navigation around simple obstacles, and still succeeds at difficult ones. While groups navigating obstacles preferentially move directly toward the nest, they change their behavior over time; the longer the ants are obstructed, the more likely they are to move away from the nest. This increases the chance of finding a path around the obstacle. Groups rapidly changed directions and rarely stalled during navigation, indicating that these ants maintain consensus even when the nest direction is blocked. While some decisions were aided by the arrival of new ants, at many key points direction changes were initiated within the group, with no apparent external cause. This ant species is highly effective at navigating complex environments, and implements a flexible strategy that works quickly for simple obstacles and still succeeds with complex obstacles.

## Introduction

From multi-cellular organization to massive animal migrations, emergent group behaviors are ubiquitous and drive much of the complexity of the biological world. Tasks are often accomplished without a leader (Camazine et al., 2001), as impressive group behavior emerges from simple individual-level interactions. Ant colonies are model systems for studying emergent group behavior due to the complexity and scale of the tasks they cooperatively accomplish. A crucial task for many animal groups, including ants, is making collective decisions, and a substantial body of studies deal with how groups accomplish this with discrete, single-step decisions (Conradt and Roper, 2005; Deneubourg and Goss, 1989; Sumpter and Pratt, 2009), such as nest site selection in honey bees or *Temnothorax* ants (Pratt, 2005; Pratt et al., 2002; Seeley, 2010). In contrast, we know less about how groups collectively accomplish complex tasks that require a series of decisions, each building on previous ones. This type of behavior is akin to problem solving, and has been studied primarily in individuals rather than groups. For example, maze-solving has been studied in many taxa including rats (Mulder et al., 2004; Yoder et al., 2011), and large, single-celled slime molds (Nakagaki et al., 2000; Reid and Beekman, 2013; Reid et al., 2012). Groups making serial decisions face the additional challenge of maintaining consensus – defined as members of a group agreeing on a single option (Sumpter and Pratt, 2009) – throughout the task. In this study we examined collective problem solving by coordinated groups of ants in a task similar to a maze.

A highly conspicuous example of collective behavior in ants is cooperative transport, in which a group of ants work together to move a large object, intact, back to their nest (reviewed in Berman et al., 2011; Czaczkes and Ratnieks, 2013; McCreery and Breed, 2014). Cooperative transport is challenging because it requires moving an object over heterogeneous terrain while maintaining consensus within the group about travel direction. Ants can generally sense the direction of their nest (e.g. Cheng et al., 2014; Steck, 2012; Wehner, 2003) and in many cases groups can accurately form consensus to move in their nest direction (Berman et al., 2011; Czaczkes et al., 2010; Gelblum et al., 2015). However, if the direction of the nest is blocked by an unexpected obstacle, the situation is substantially more challenging. The shared homeward bias of the group is no longer helpful. In order to proceed the group must find a consensus on a new travel direction and navigate around the obstacle, continuously updating the travel direction until it is possible to resume unobstructed movement toward the nest.

How a group solves this problem – their *strategy* – impacts the kinds of obstacles they can successfully navigate. Consider a simple strategy: when a transport group encounters an obstacle, they choose a direction to move around obstacle perimeter until it is possible again to move, unobstructed, toward the nest. This strategy requires information about nest direction and the ability to form consensus on travel direction, both of which are plausible for ant colonies. However, this simple strategy only works with simple obstacles; groups using this strategy would get stuck in even a slightly complex obstacle with a concave shape, which would require moving away from the nest to succeed. On the other hand, navigation strategies exist that would be successful for any possible obstacle, but require more information and updating (e.g. Table 1; Kamon and Rivlin, 1997; Murphy, 2000). We define these strategies as “robust” because they are successful over a broad range of conditions. But robustness comes at a cost in terms of energy and information processing. Thus, there is a trade-off between simple strategies that are easy for groups to execute but may fail, and robust strategies that are more costly.

We investigate obstacle navigation during cooperative transport in *Paratrechina longicornis* (Latreille), the longhorn crazy ant. Workers of this species are known to be excellent transporters (Czaczkes et al., 2013; Gelblum et al., 2015). *P. longicornis* is in the subfamily Formicinae and is a widely distributed “tramp” ant (Wetterer, 2008). We presented groups of ants with obstacles of varying difficulty in order to look at the navigation strategy they use, and where their strategy falls between simple and robust problem solving. We obstructed ant cooperative transport groups with three obstacles of increasing complexity: an obstacle that simple strategies can easily navigate (the “wall”), an obstacle that requires a more robust strategy (the “cul-de-sac”), and an impossible obstacle that thwarts even robust strategies (the “trap”) (Fig. 1). Example strategies with their predictions are shown in Table 1. Our main questions were: 1) How robust is the strategy of groups of ants? 2) What strategy do the ants use? 3) How do individuals contribute to the group's strategy? 4) When facing an obstacle that is impossible to navigate, do groups have the ability to detect traps, and if so what is their response?

**Fig. 1:**
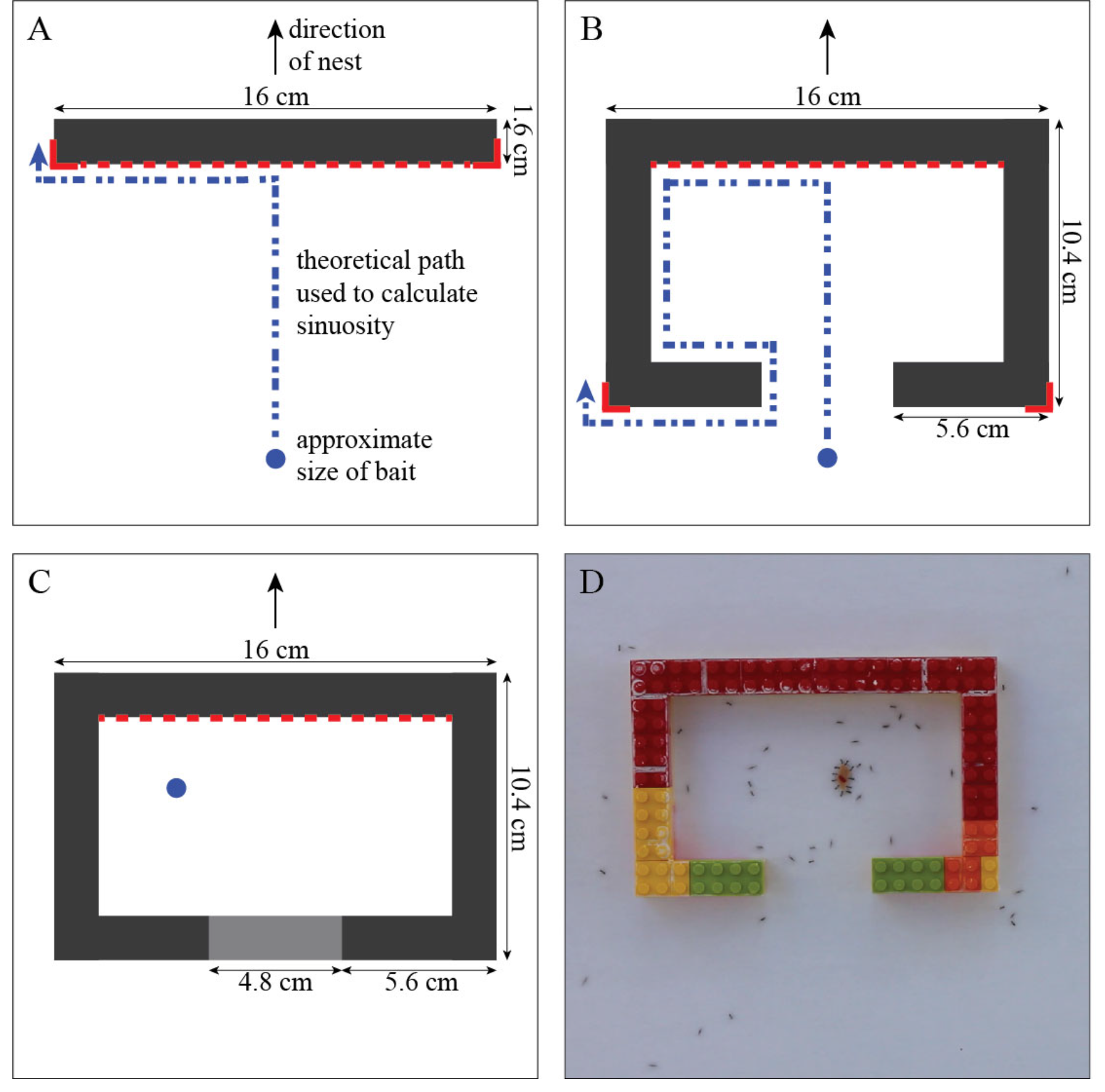
Shapes and dimensions of obstacles used to block cooperative transport groups. A: The wall; B: the cul-de-sac; and c: the trap. Red dashed line indicates the “back wall” as referenced in the text. We defined obstacle navigation as beginning when the group reached this back wall, and ending when the group had rounded one of the corners marked in red. The blue dashed line indicates the theoretical “shortest” path used to calculate sinuosity, and the blue circle indicates the approximate size of the bait. D: An example of a trial with the cul-de-sac.

**Table 1:** Predictions for efficiency of navigation in the wall and the cul-de-sac with example strategies of different robustness.

| Example Strategy | Description | Prediction |
| --- | --- | --- |
| None | Groups move in the direction of the nest, however when that direction is obstructed they are unable to form consensus on a new direction. | Groups fail to navigate any of the obstacles. |
| Simple 1 | Groups move in the direction of the nest whenever possible. If obstructed, the group can form consensus on a new direction, and they follow the obstacle perimeter until the nest direction is available. This requires the ability estimate nest direction and to form consensus on a travel direction even when the nest direction is unavailable. | Groups succeed at navigating the wall but fail to navigate the cul-de-sac, because in the cul-de-sac the nest direction becomes available before navigation is complete. This strategy fails on all non-convex obstacles (Lumelsky and Stepanov, 1987). |
| Extremely robust 1 | Groups move towards nest direction whenever possible; if obstructed, groups choose move around obstacle as in the Simple 1. However, the group only leaves the obstacle perimeter when the nest direction is available and the nest is closer than at any other point during navigation. This strategy requires the ability to estimate distance to the nest. | Groups will successfully navigate both the wall and the cul-de-sac with maximum efficiency (sinuosity = 1 for each). This strategy is known to solve arbitrarily complex obstacles (Kamon and Rivlin, 1997). Multiple strategies exist that are extremely robust, but all require more information processing than Simple 1. |
| Extremely robust 2 | Groups follow the “extremely robust 1” described above, but if they navigate over ground they have already moved over, they abandon the transport effort. This requires detecting when they move over their footsteps, which can be accomplished through path integration, for example. | Groups will successfully navigate both the wall and the cul-de-sac with sinuosity = 1 for each. This strategy also succeeds in the trap because individuals can give up, moving on to other useful behaviors. |
| Intermediate strategies | There are many possible strategies that would lead to moderately robust navigation. All require more information than the simplest strategy. Using trajectory data, we can infer properties of the strategy. | Groups succeed at both the wall and the cul-de-sac. Efficiency for the wall may be higher (lower sinuosity) than for the cul-de-sac. Efficiency is lower than an extremely robust strategy (sinuosity > 1) for both obstacles. |

## Methods

### Overview

We gave ants foraging near a colony entrance pieces of tuna to carry. After a group of ants had begun carrying the tuna, and their preferred direction of travel was established, we put one of three obstacles in their path, directly blocking that preferred direction. We video recorded these trials and extracted data from the videos, including the trajectory of the piece of tuna, which we used to measure additional results metrics, described below.

For question 1, we examined strategy robustness by looking at which obstacles groups of ants could navigate, their efficiency at the wall and the cul-de-sac, and how well they maintained consensus about travel direction (Table 1). For question 2, we identified behavioral elements (e.g. perimeter following, direction changes) that make up the strategy. For question 3, we examined individual behaviors during key decisions in obstacle navigation, to look for precipitating events such as ants joining. For question 4 we compared group behavior in the cul-de-sac and the trap.

### Study sites

We conducted fieldwork in June 2014 at two sites: one at the Arizona State University in Tempe, Arizona, and the other at the Biosphere 2 facility in Oracle, Arizona. We conducted experiments at two colonies in Tempe and two at the Biosphere 2 facility (4 colonies total). We worked in locations fitting having a flat surface, with relatively constant shade, and close to only one nest entrance so that all foragers had the same goal. These constraints limited the number of colonies available to us. Nevertheless, given the variation in behavior that we observed, four colonies is sufficient to answer our questions. Colonies varied in traits such as group size and speed, as discussed below, but the general pattern of navigation behavior was similar among these colonies.

### Obstacles and strategy

We used three types of obstacles. 1) The simplest obstacle (Fig. 1A), hereafter referred to as the “wall,” requires a form of symmetry breaking: groups must choose a direction from equal options. We placed obstacles approximately perpendicular to groups' travel direction, blocking the nest direction. Two of the remaining options (left and right along the perimeter) are of equal value, so a choice between them requires breaking symmetry. 2) A complex obstacle (Fig. 1B), the “cul-de-sac,” also requires symmetry-breaking when first encountered, but has an additional challenge. Groups navigating this obstacle must move opposite to their preferred direction (away from the nest) to succeed. The last obstacle (Fig. 1C), the “trap,” resembles the cul-de-sac but is impossible to navigate. However there are strategies that would at least allow groups to know that they are trapped. In natural settings, such strategies would be more robust, because if groups recognize that navigation has failed, individual ants can abandon the transport effort and switch to other useful behaviors. For example, without their load, ants may be able to escape and return to the nest.

Navigation strategies are also of interest in robot navigation, and a substantial body of literature predicts the consequences of various navigation strategies in the presence of obstacles of varying complexity (Kamon and Rivlin, 1997; Lumelsky and Stepanov, 1987; Murphy, 2000). While we do not expect ants to use any specific strategy from this literature, we used these predictions to design our obstacles, so that we know a group level strategy that can solve each one. We can compare the ant groups' trajectories to the predictions for these theoretical strategies, listed in Table 1. Strategies that succeed with a wider range of obstacle types require more information processing than simpler strategies. The strategies included in Table 1 are just examples; the range of potential strategies is large, and strategies could include more stochasticity. For example, groups may follow the perimeter of an obstacle until the nest direction is available, at which point they move toward the nest with a certain probability, otherwise continuing to follow the perimeter. Stochastic strategies may allow groups to navigate a wider range of obstacles, but their efficiency with a given obstacle will be different in each encounter.

All potential strategies are composed of behavioral elements. Examples of elements included in the strategies in Table 1 are perimeter following, moving toward the nest, moving away from the nest, and remembering the path travelled. Other possible elements include spontaneous direction changes and random walks.

### Experiment details

At the beginning of each trial, a fresh piece of 11×17 inch (28×43 cm) white paper was placed on a flat surface near a nest entrance. We set up in locations where all successful foragers went back to the same entrance, to ensure that individuals in our transport groups would have the same goal. The nest entrance was at least 15 cm away from our experiment. At the start of each trial a dead cricket was placed on the paper, so that foragers would recruit by laying pheromone. We used a cricket to elicit a strong recruitment response, so that transports would not fail because of insufficient workers. When a group of workers began moving the cricket, we replaced it with a marked piece of tuna, lighter than the cricket (0.031 g - 0.105 g). Once a group of workers had moved the tuna at least 10 cm, one of the obstacles was placed in their path, oriented such that the “back wall” (dashed red line in Fig. 1) was perpendicular to their preferred direction. For the trap, we first obstructed their path with an obstacle shaped like the cul-de-sac. After the group entered this obstacle, we placed a “door” in the exit so that groups could not leave. We did not try to eliminate additional ants from being in the vicinity of the transport effort. These “extra” ants, also known as escorts, are common in natural *P. longicornis* transport efforts (Czaczkes et al., 2013), and whatever affect they have on navigation would also be present in natural navigation efforts.

These escort ants did not alter pheromone trails to help groups navigate around obstacles. *P. longicornis* workers lay a specialized pheromone trail to recruit ants to a large item, but cooperative transport groups do not use that trail to navigate back to the nest (Gelblum et al., 2015). The recruitment trail is short-lived, decaying within 6 minutes (Czaczkes et al., 2013), and workers laying this trail have a conspicuous, halting movement pattern which we did not observe during navigation. Instead of relying on a pheromone trail, recent studies suggest that transport groups in *P. longicornis* can be aided in returning to the nest by new ants joining the effort (Gelblum et al., 2015). To rule out the possibility that other workers provide another type of global directional cue, we conducted an experiment to see how long an obstacle must be in place before ants avoid it altogether. In our experiment, the vast majority of ants moving from sugar-water baits back toward a nest initially hit the obstacle we placed in their path, but eventually ants' paths changed so that they avoided the obstacle. However, this change took more than 20 minutes, while our longest trial was only 10.8 minutes (Fig. S1). Over the length of time that our trials lasted, ants' paths had not substantially changed to avoid obstacles. Escort ants may yet provide cues to cooperative transport groups through physical contact or another mechanism, but it is unlikely that they provide global cues observable by the entire group.

All trials were recorded using a Canon Rebel T2i with lens EF-S 18-55IS (1920×1080, 30 frames per second; Canon, Tokyo, Japan). All obstacles were constructed out of Lego^®^ (Billund, Denmark) and coated on the inside with Insect-a-Slip (Fluon; BioQuip, Gardena, CA, U.S.A.) so that groups could not climb over them.

A total of 91 trials were conducted across the four colonies. However, we excluded trials in which the following occurred: 1) foragers recruited from multiple nest entrances, 2) the tuna piece was light enough to be moved substantial distances by a single ant, or 3) the group first encountered the obstacle either by hitting a side wall as opposed to the back wall, or by hitting the back wall but more than 3.2 cm away from the center. After excluding these trials we were left with a total of 61 trials: 22 trials for the wall, 19 trials for the cul-de-sac, and 20 trials for the trap.

### Data extraction

Several types of data were extracted from each of these 61 videos. We manually recorded the location and orientation of the tuna piece every second (30 frames) using Matlab. This provides the trajectory of the group rather than of individual ants. The location of the obstacle with respect to each trajectory was also recorded. We used this trajectory information to measure speed, sinuosity, and backward runs, and to identify the sharpest turns in the wall and escape points in the cul-de-sac, as described below.

*Speed (questions 1, 3, and 4):* Speed was measured every second, so that it approximates instantaneous speed. At times when the speed was extremely low (less than 0.048 cm s^-1^) we classified the group as being “stalled.” For analyses in which we compared speeds across trials, we eliminated an initial period when the group was getting started. To do this, we removed speed data for early times until the group had first reached a threshold speed of at least 0.24 cm s^-1^.

*Sinuosity (question 1):* Sinuosity is defined as the ratio of the path length to length of the shortest possible path. Paths with lower sinuosity are more efficient. Here, in order to directly compare obstacles, we used a modified measure of sinuosity: the path length divided by the path that would be taken if groups followed the perimeter of the obstacle (Fig. 1A,B).

*Backward runs (questions 2 and 4):* As discussed above, for the cul-de-sac, successful navigation requires moving away from the nest. To see the extent to which groups moved opposite to the preferred direction, we quantified the number and distance of “backward runs.” A backwards run is a period of time during which the group is moving away from the nest (and away from the back wall). Backwards runs occur when the distance from the piece of tuna to the back wall is increasing, and the run ends as soon as this distance decreases. For each backwards run we recorded the time at which the run started, and the total displacement away from the back wall that occurred during that run.

*Sharp turns (question 3):* Transport groups sometimes turn sharply during navigation. In these instances the consensus travel direction changes. We carefully examined these sharp turns to determine what may have caused these changes in consensus. To simplify our analysis we focused only on sharp turns occurring while groups navigated the wall. For each trial we calculated a turning angle every second by taking the mean direction over the three previous seconds and over the three subsequent seconds. We carefully examined every point that had a turning angle equal to or greater than 2.5 radians (~145 degrees). We chose 2.5 radians because it resulted in 38 unique turns for all the wall trials, which was a reasonable number to examine manually. Each of these turns was carefully examined and placed into one of the following four categories based on what may have caused it: 1) a new ant joined the transport effort, 2) an ant left the transport effort, 3) the group hit the obstacle, and 4) there was no discernable cause (i.e. none of the above).

*Escape points (question 3):* Escape points are turns that led to successful completion of the navigation. We define the escape point of each trial as the last turn the group made, such that after making the turn they left the interior of the cul-de-sac, while before making the turn they would not have done so. We manually determined the escape point for each cul-de-sac trial from images of the complete trajectories; our designation of escape points was therefore blind with respect to the behaviour of individuals. We placed each escape point into one of the same four categories listed for sharp turns.

In addition to these results metrics that we measured from group trajectories, we manually measured navigation time (for question 1) and group size (for question 3). For navigation time, we defined the start of navigation as when a group first appeared to respond to the “back wall,” and we defined the end of navigation as when they rounded one of the bottom corners (colored in solid red in Fig. 1) and could move unobstructed towards the nest. To find group size we counted the number of ants attached to the tuna every 15 seconds (every 450 frames).

### Statistical Analysis

All statistical analyses were performed in R, version 3.2.2 (R Core Team, 2015).

*Question 1:* We compared efficiency data in the wall and the cul-de-sac with a t-test on sinuosities, log-transformed to meet assumptions of normality. To look at how well groups maintain consensus, we compared stalls and speeds while groups were navigating and while unobstructed. The proportion of time groups were stalled was analyzed with a Bayesian zero-inflated beta model using Stan implemented in R (Stan Development Team, 2015). We chose this method because these data were heavily zero-inflated, such that general or generalized linear models were inappropriate. We used a binomial model to look at the probability of never stalling, and for trials with at least one stall, we used a beta model to look at the proportion of time stalled. We looked at the effect of potential predictors on this proportion, but not on the probability of never stalling, because this probability is biased among obstacles: groups are less likely to never stall in the cul-de-sac simply because it takes them longer to navigate it. Priors were relatively vague and did not substantially impact posteriors.

We used general linear models to analyze square-root transformed speeds using the lme4 package (Bates et al., 2015, 4). Transformed speeds were normally distributed. As potential predictor variables, we included whether the group was navigating an obstacle or unobstructed, and which obstacle was navigated. We compared a full model with both predictors and their interaction to simpler models with one or both predictors without the interaction (Table S1). We also included random effects of trial nested within colony. We used a likelihood approach, comparing models with Akaike Information Criteria (AIC) to determine the best predictor variable(s).

*Question 2:* The distances of backward runs were analyzed in a Bayesian framework using JAGS in R (Plummer et al., 2015). We saw how behavior changed over time by modeling the distribution of backward runs distances as a gamma distribution, where the shape parameter, k, can change in time according to the following equation.

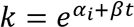

Where *α_i_* represents a random intercept for each trial and *β* indicates the extent to which k changes over time. Gamma distributions with larger shape parameters are more right-skewed.We also fit additional parameters indicating the scale of the beta distribution and the mean and standard deviation of *α_i_*. We verified this model by simulating hundreds of data sets and checking that the 95% credible interval included the true parameter 95% of the time. We looked at whether the distribution of backward run distances changes over time by evaluating whether *β* is different from zero. Our priors did not substantially impact posteriors. To avoid bias in the timing of large backwards runs, we removed from the analysis any backward run that led to the group completing obstacle navigation.

*Question 3:* In addition to a qualitative analysis of sharp turns and escape points, we examined the effects of group size and speed on performance using Pearson product-moment correlations, with log-transformed data where necessary. We also looked at the effect of number of ants on speed using a Kendall rank correlation, as speed data could not be transformed appropriately for a Pearson correlation due to non-normality.

*Question 4:* We qualitatively analyzed speeds over time in the trap and the cul-de-sac.We used the ggplot2 package (Wickham, 2009) to visually compare these speeds, smoothing the speed data with local regression (LOESS) or generalized additive models (GAM), depending on the number of observations. We also compared stalls and backwards runs in the cul-de-sac and the trap, using the statistical tests described above.

## Results

### 1) *How robust is the navigation strategy of groups of ants?*

*P. longicornis* groups successfully navigated both the wall (n = 22) and the cul-de-sac (n = 19) in every trial. As expected, they never successfully navigated the trap (n = 20). Examples of groups' trajectories are shown in Fig. 2 (Movie S1 shows the complete navigation for Fig. 2A,C; Figs. S2, S3, and S4 show group trajectories for all trials with the wall, cul-de-sac, and trap, respectively). The mean times to navigate the obstacles were 1.01 minutes *(s.e.m.* = 0.17) for the wall and 5.99 minutes *(s.e.m.* = 0.52) for the cul-de-sac, with a mean speed across trials of 0.43 cm s^-1^. While groups always solved both obstacles, they were significantly more efficient when solving the wall than the cul-de-sac, (untransformed sinuosity means: 2.5 *(s.e.m.* 0.36) and 6.3 *(s.e.m.* 0.80); unpaired *t* test, t_39_ = 5.05, *P* < 0.0001; Fig. 3).

We also looked at how well groups maintain consensus about direction throughout the navigation process in the wall and the cul-de-sac. We expect groups lacking consensus to stall. Transport groups rarely stalled (< 2% of the time) either while navigating obstacles or while unobstructed (Table 2). Groups had a 54% chance of never stalling (probability = 0.54, 95% credible interval (CI): 0.43 – 0.64). For trials in which at least one stall occurred, groups spent the same proportion of time stalled regardless of whether they were obstructed. Posterior distributions overlapped substantially for unobstructed groups (mean = 0.044, 95% CI: 0.025 – 0.081), groups navigating the wall (mean = 0.044, 95% CI: 0.021 – 0.084), and groups navigating the cul-de-sac (mean = 0.025, 95% CI: 0.014 – 0.052). We estimated random intercepts for each colony, but found that colonies did not differ in proportion of time stalled (Fig. S5). The results show that groups rarely stall, and the proportion of time groups are stalled is not affected by the presence of either obstacle.

**Table 2:** Proportion of time spent stalled while unobstructed (before or after obstacle navigation) and during obstacle navigation. Values shown are means of these proportions for each trial.

| Obstacle | Mean proportion of time stalled<br>(including trials with zero stalls) |  |
| --- | --- | --- |
|  | *While unobstructed* | *While navigating obstacle* |
| Wall ($n = 22$) | 0.014 | 0.016 |
| Cul-de-sac ($n = 19$) | 0.017 | 0.016 |
| Trap ($n = 20$) | 0.014 | 0.24 |

**Fig. 2:**
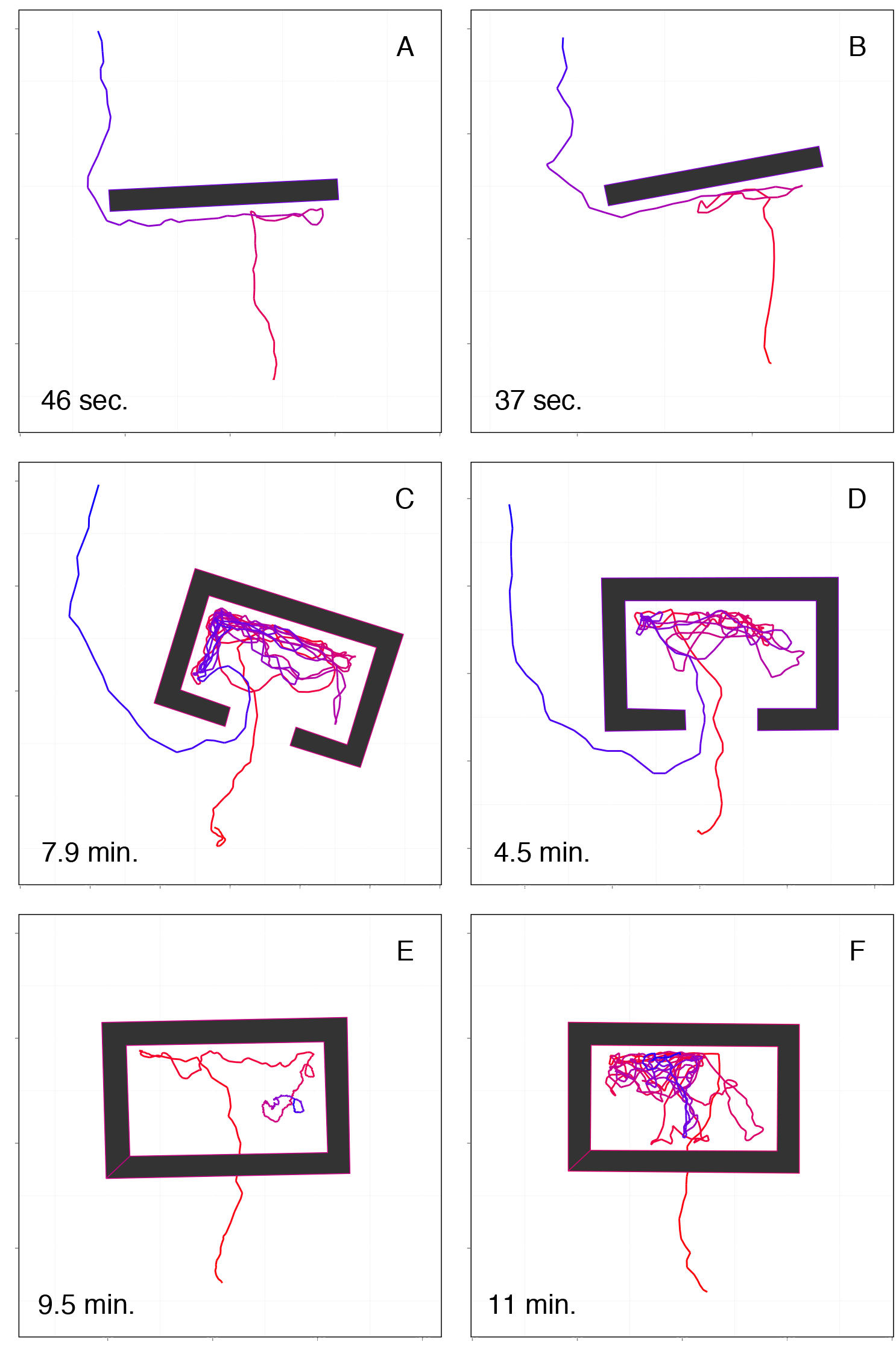
Examples of group trajectories. Warmer colors indicate early points in the navigation process, while cooler colors are later in time. A, B: the wall; C, D: the cul-de-sac; E, F: the trap. Times shown in the bottom right corner of each panel indicate the time it took to navigate the obstacle (from hitting the back wall to rounding the corner) for A through D, and the time spent trapped for E and F.

**Fig. 3:**
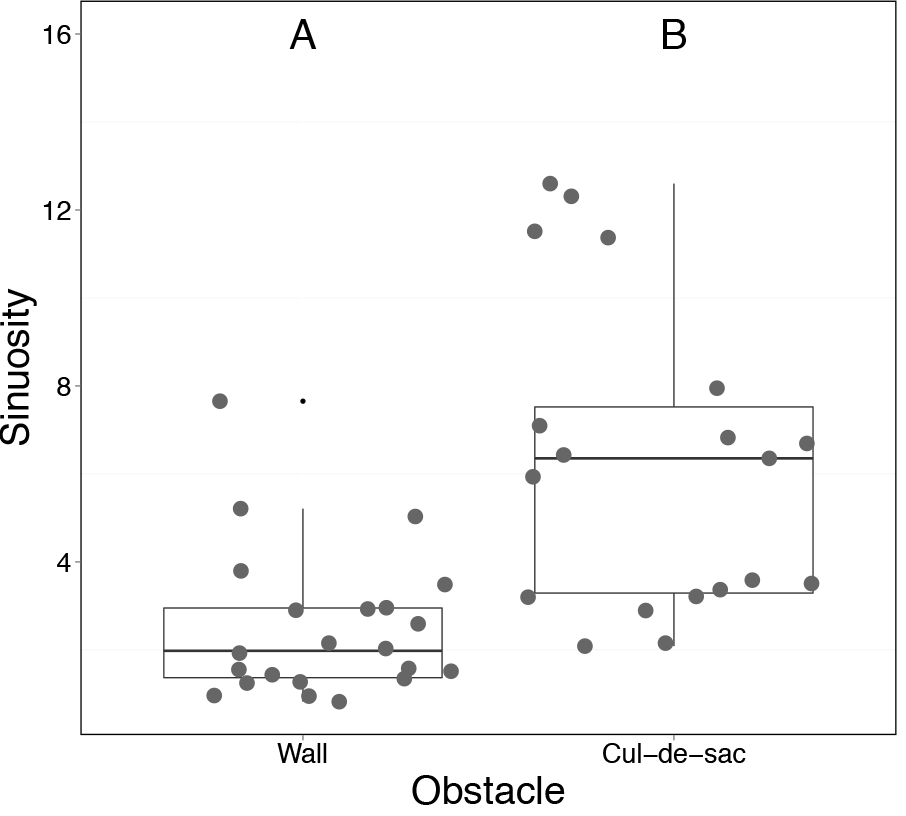
Efficiency (sinuosity) of cooperative transport for groups navigating the wall (n = 22) and culde-sac (n = 19). Unpaired t test, *t*_39_ = 5.05, p < 0.0001. Large dots show jittered sinuosity values for each trial. Letters indicate groups are significantly different. Boxes include 50% of the data (going from the 25th to 75th percentiles), and whiskers extend to the lowest and highest values that are within 150% of the interquartile range. Small dots are points outside that range.

To look further at consensus, we evaluated the effect of the obstacle on speed using general linear models. The best model included whether the group was obstructed (*β* = -0.065), and did not include whether they were navigating the wall or the cul-de-sac. Detailed results of this best-fit model are included in Table S2. Groups slow down while navigating obstacles and this effect is not different for different types of obstacles. However, the effect size was small; all else being equal, groups moved only 0.043 cm s^-1^ slower while navigating obstacles. This is approximately a 10% reduction in speed, and this decrease is much smaller than the overall variation in speeds of groups (Fig. 4).

**Fig. 4:**
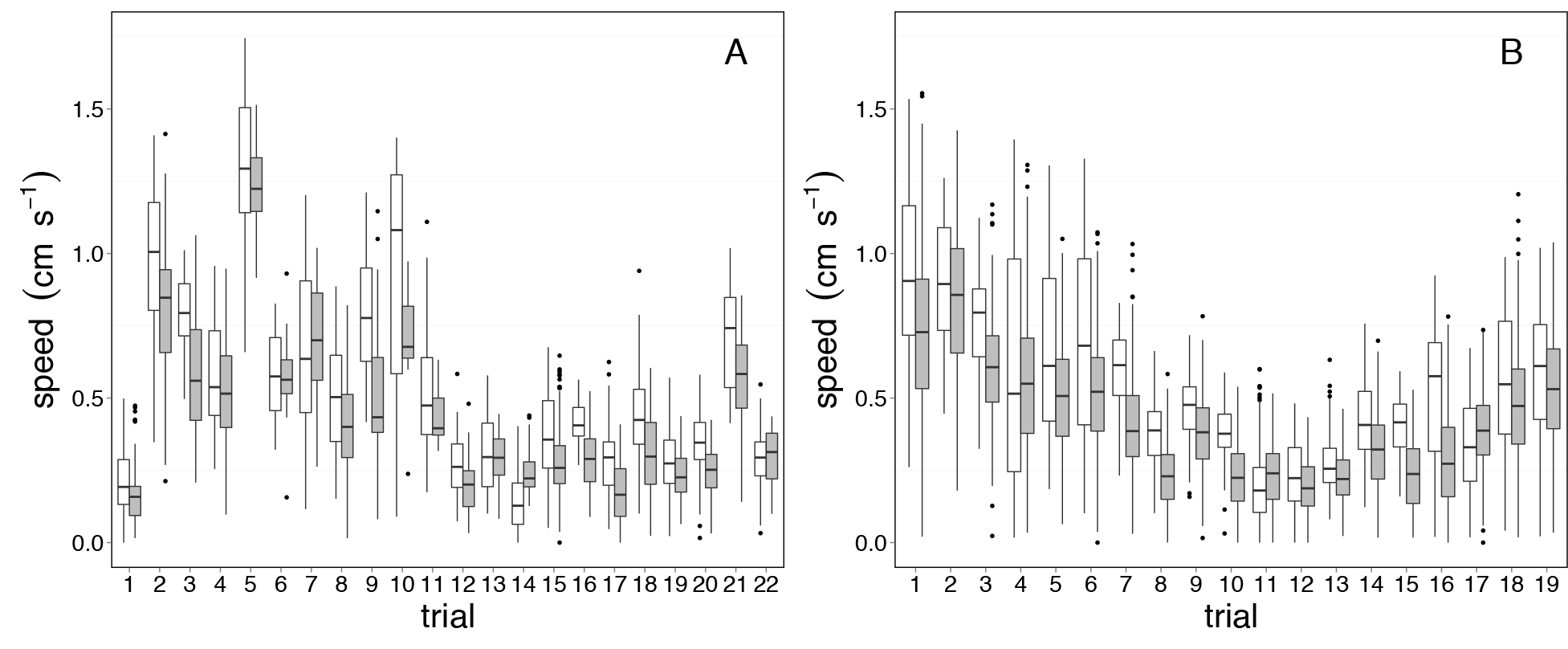
Mean speeds for groups navigating an obstacle (gray boxes) and while unobstructed (open boxes) for each trial. A: trials with the wall; B: trials with the cul-de-sac. Boxes include 50% of the data (going from the 25th to 75th percentiles), and whiskers extend to the lowest and highest values that are within 150% of the interquartile range. Dots are points outside that range. The best general linear model of speeds, determined using AIC, included whether the group was obstructed (*β* = -0.065) and a random effect of trial nested within colony

### 2) *What strategy do groups use?*

We looked at elements – the behavioral components of strategy – to get at this question. The primary challenge in navigating the wall is to break symmetry. We could not ensure perfect symmetry among the options for travel direction (obstacles were not always perfectly perpendicular to preferred direction), but groups did not favor the direction closer to their initial travel direction. Upon hitting an obstacle, *P. longicornis* groups broke symmetry and chose a single direction to move around the wall. They did not, however, stick with that chosen direction for the entire navigation. In 14 out of 22 trials (64%), the group changed direction after the initial choice. The groups also navigated directly up to the wall, and then typically remained fairly close to the wall perimeter after choosing a travel direction (Figs. 2 and S2). There was no evidence of visual detection and evasion; transport groups did not start avoiding any of the obstacles from a distance.

The cul-de-sac presents an additional challenge in that groups must move counter to their preferred direction to successfully navigate this obstacle. In this case initial behaviors of groups were similar to those of groups navigating the wall; transport groups first hit the back wall of the obstacle and quickly picked a direction. Early in the navigation process, groups typically remained close to the back wall, but they did not follow the perimeter and move along the side walls (Fig. 2). Later in the navigation process groups moved away from the perimeter into the open area within the cul-de-sac. Our results show that groups moved further backwards (away from the nest) the longer they had been navigating the cul-de-sac (Fig. 5B). More specifically, the distribution of distances of backwards runs becomes more right-skewed (shifted toward longer distances backwards) over time. Our estimate of *β* in the cul-de-sac is 0.13, and the 95% credible interval (CI) is 0.07 – 0.20. We did not find strongevidence for this effect in the wall (*β* = 0.0019, 95% CI: -0.15 – 0.15, Fig. 5A), perhaps because groups navigated the wall rapidly. Transport groups can change their behavior over time. Estimates for all parameters for this analysis are in Table S3.

**Fig. 5:**
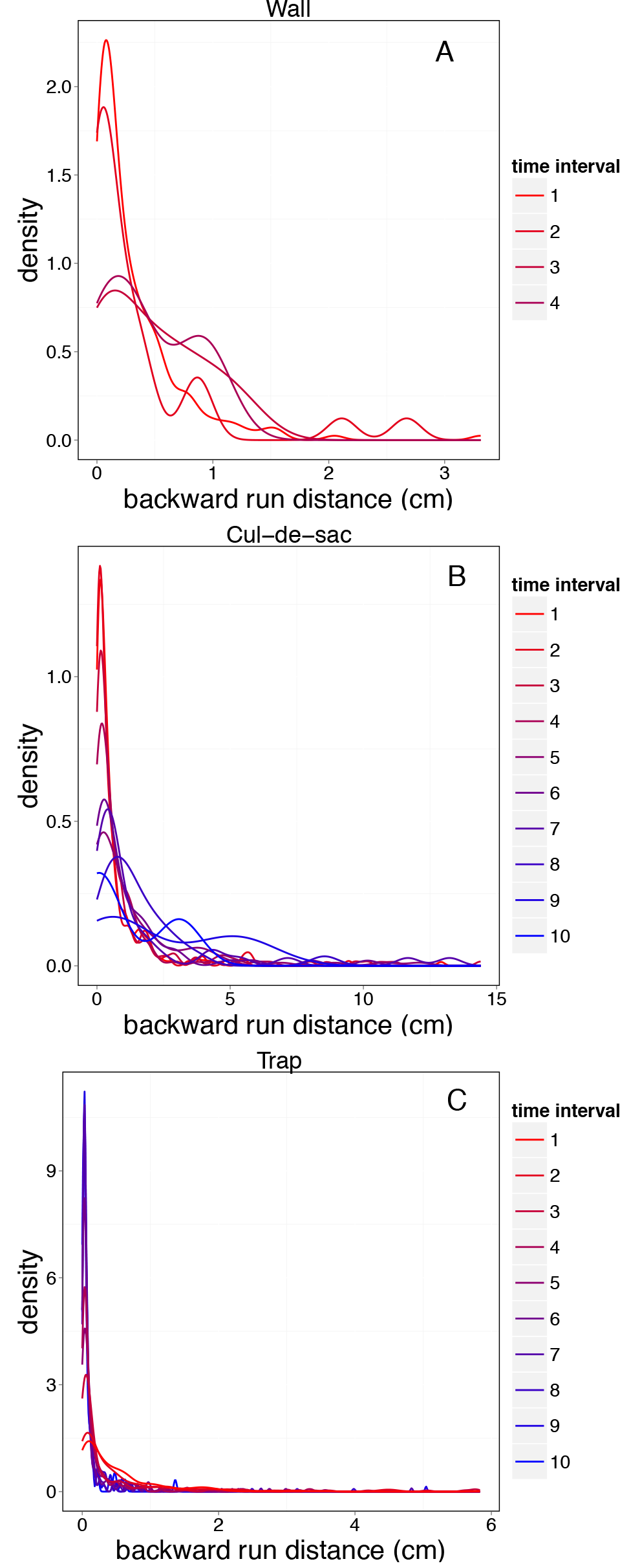
Densities of backward run distances at different time intervals. A: the wall; B: the cul-de-sac; and C: the trap. Warmer colors indicate earlier time intervals. Modeled in a Bayesian framework as a gamma distribution with a changing shape parameter, our estimate of the effect of time, β, in the cul-de-sac (B), is 0.13 (05% CI: 0.07 – 0.20). We did not find strong evidence for this effect in the wall (A; *β* = 0.0019, 95% CI: -0.15 – 0.15), and found the opposite effect in the trap (C; *β* = -0.22, 95% CI: -0.26 – -0.19).

### 3) *How do individuals contribute to the group's strategy?*

We observed many sharp turns during these experiments, yet as discussed above, stalls were very rare. Thus, groups were able to sharply change travel direction without stalling. To investigate what happens during those turns, we qualitatively examined the sharpest turns made while navigating the wall (n = 38). In 32% of cases the change in direction appeared to be caused by a new ant joining the transport group, in 24% of cases the group hit the obstacle, and in 45% of cases we could see no precipitating event in the moments preceding the direction change (Table 3). We conducted a similar analysis of escape points in the cul-de-sac (Table 3). In over half of cases (58%) we could detect no events that seemed to cause the direction changes leading to escape, while 32% were caused by new ants.

**Table 3:** Apparent reasons for the sharpest turns during wall navigations (n = 38), and for the last turn leading to escape from the cul-de-sac (n = 19).

|  | **Ant**<br>**joined** | **Ant left** | **Hit**<br>**obstacle** | **No discernable**<br>**reason** |
| --- | --- | --- | --- | --- |
| *Sharp turns* |  |  |  |  |
| **Number**<br>(out of 38) | 12 | 0 | 9 | 17 |
| **Percent** | 32% | 0% | 24% | 45% |
| *Escape points* |  |  |  |  |
| **Number**<br>(out of 19) | 6 | 2 | 0 | 11 |
| **Percent** | 32% | 11% | 0% | 58% |

In addition to individual contributions at key moments, we looked at what effect the number of ants in a group had on transport efficiency. Within the range of group sizes we observed, transport groups with more ants were faster: Number of ants and speed were correlated through time (Kendall's *τ* = 0.44, *P* < 0.0001). This correlation was present within colonies as well as in the pooled data (Fig. S6). However, groups navigating the wall with higher average speeds were not more efficient with respect to sinuosity (Pearson's *r* = – 0.14, *P* = 0.53; Fig. 6A). Furthermore, for the cul-de-sac, faster groups were *less* efficient, in that they had higher sinuosity (Pearson's *r* = 0.62, *P* < 0.01; Fig. 6B). This correlation in the cul-de-sac between speed and sinuosity was present across colonies, but not within individual colonies. Colonies at Arizona State University tended to have higher speeds and sinuosities, while colonies at Biosphere 2 tended to have lower speeds and sinuosities (Fig. 6B). Groups with more ants, on average, while navigating obstacles were not more efficient with respect to sinuosity (wall: Pearson's *r* = −0.18, *P* = 0.43, cul-de-sac: Pearson's *r* = 0.25, *P* = 0.30, Fig. S7).

**Fig. 6:**
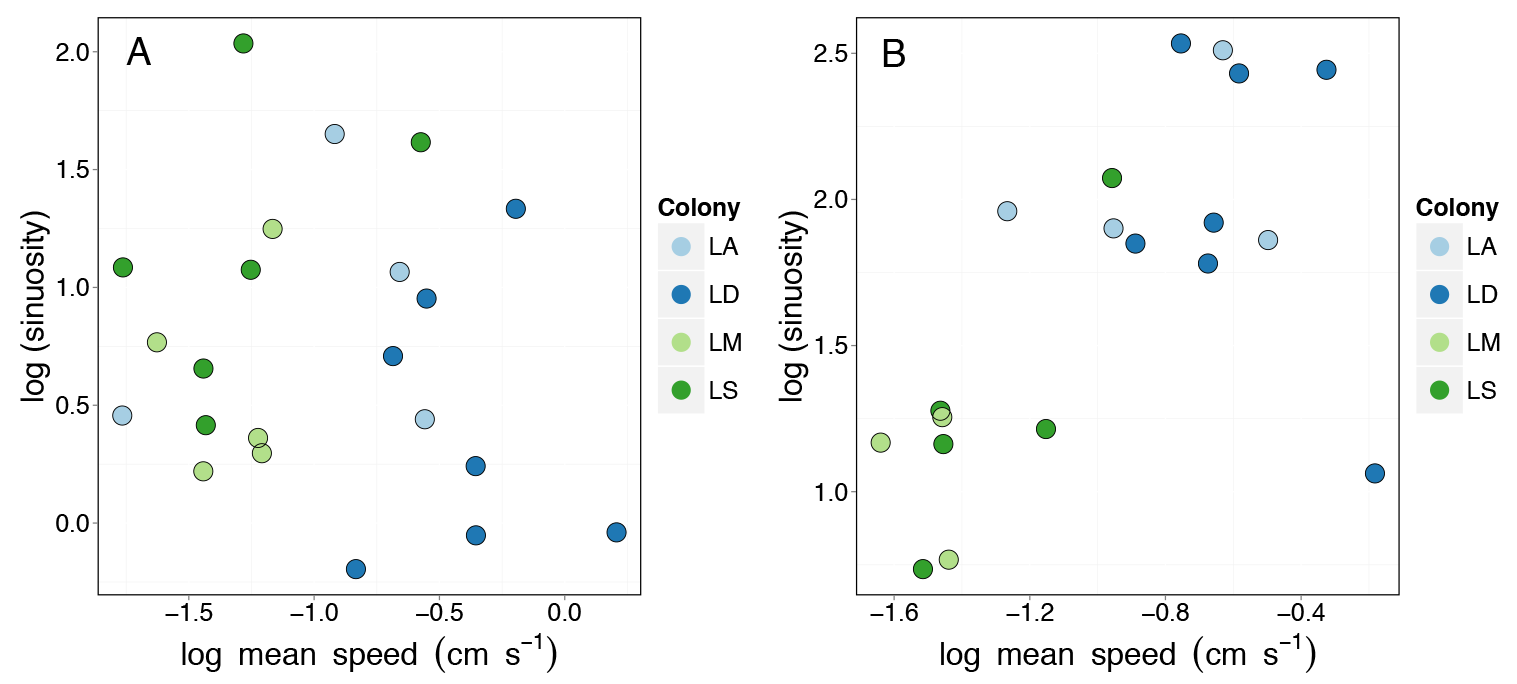
Mean speed of groups is not correlated with sinuosity for groups navigating the wall (A; Pearson's *r* = -0.14, *P* = 0.53), and is positively correlated with sinuosity for groups navigating the cul-de-sac (B; Pearson's *r* = 0.62, *P* < 0.01). Blue dots are points for colonies at Arizona State University (light blue: colony LA; dark blue: colony LD) and green dots indicate colonies

### 4) *How do groups behave with an impossible obstacle?*

The aggregate behavior of groups in the trap was starkly different from group behavior in either the wall or the cul-de-sac, across multiple results metrics. We expected groups to behave similarly in the trap and the cul-de-sac at least initially, however the speeds of groups in these obstacles differ even early on in the navigation process, with groups in the trap slowing down dramatically (Fig. 7). This drop in speed corresponds to an order-of-magnitude increase in the amount of time groups are stalled (Table 2). Furthermore, while our analysis of backwards runs indicates that groups have larger backwards runs the longer they have been in the cul-de-sac, we found the opposite pattern for the trap *(β* = -0.22, 95% CI: -0.26 – -0.19, Fig. 5C). This pattern likely results from groups simply moving more slowly and less. In the trap, over time, groups explore the space less and less.

**Fig. 7:**
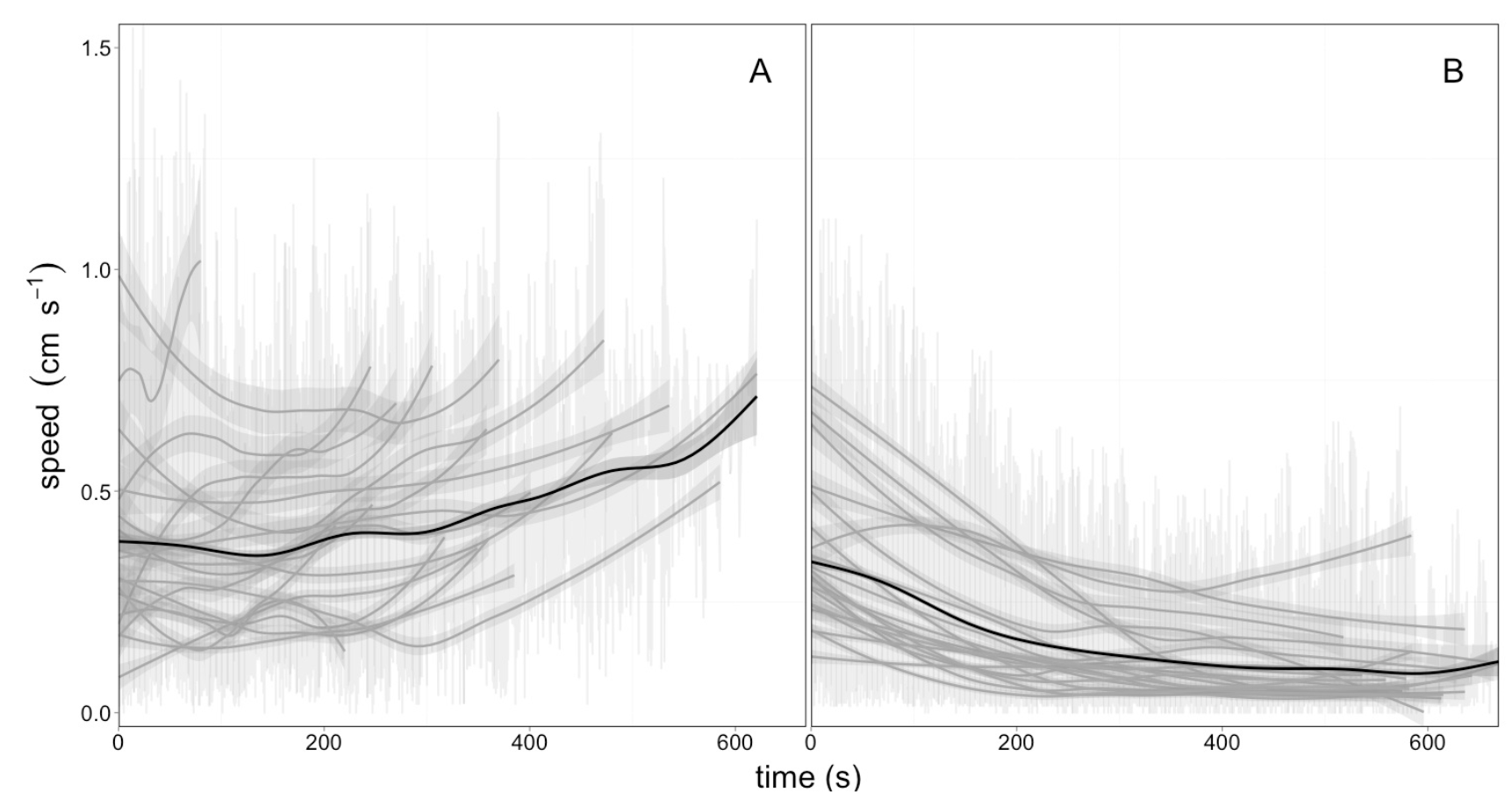
Speeds of groups over time navigating the cul-de-sac (A) and in the trap (B). Groups in the trap reduce their speed dramatically, while groups in the cul-de-sac maintain relatively constant speeds. Light grey, unsmoothed lines (background) show raw speed data. Grey, smooth lines show the smoothed speed for each trial, computed with LOESS, and black lines show smoothed speed across trials, computed with GAM.

## Discussion

We presented *P. longicornis* cooperative transport groups with obstacles that require a series of decisions to navigate (problem solving). From their responses we infer what kind of strategy they use. We predicted a simple strategy based on nest direction would fail at the cul-de-sac, while an extremely robust strategy would be maximally efficient (sinuosity = 1) for both the wall and the cul-de-sac. Neither of these extreme results occurred. Instead, *P. longicornis* uses a moderately robust strategy. Groups of ants were more efficient – took more direct paths in less time – when navigating the wall than the cul-de-sac, but groups still always succeeded in solving the cul-de-sac. Furthermore, their strategy was stochastic. Trajectories in the same obstacle differed greatly from one trial to the next, and the groups did not respond predictably when facing a given set of circumstances. While we do not know exactly how robust their strategy is, we show that it is *more* robust than one based only on nest direction. In terms of the trade-off between simple, inexpensive strategies that may fail and robust strategies that require more information, this ant species appears to have a solution that lies in between. They can navigate complex obstacles, but not without some cost in terms of efficiency.

In addition to showing that groups can navigate complex obstacles, specifically the cul-de-sac, we were also able to determine *how* they succeed. Groups have stochastic behavior that changes over time. Early after encountering an obstacle, groups were unlikely to move substantial distances “backwards” (away from the nest). This is consistent with groups initially using a simple strategy in which they move in the direction of the nest whenever possible. This strategy was successful with the wall but not with the cul-de-sac. Groups navigating the cul-de-sac gradually changed their behavior by incorporating longer backwards movements, allowing them to find the exit. If groups had incorporated many backwards runs from the beginning, it would have taken them longer to navigate an obstacle like the wall. Thus, changing their strategy over time allows them to rapidly solve simple obstacles, while still eventually succeeding at complex ones.

Regardless of the specific obstacle being navigated, groups were highly effective at maintaining consensus while making a large number of serial decisions. These groups chose an initial travel direction after coming up against the obstacle, and also frequently decided to change direction. They decided to move backwards or they decided to try moving toward the nest again. When groups changed direction, they did so rapidly, without stalling, as the proportion of time stalled was the same whether or not they were obstructed. Stalls were also rare unless they were trapped (Table 2), and their speed was only slightly reduced while navigating obstacles. In the absence of obstacles, maintaining consensus is easier because groups simply move in the direction of the nest. We conclude that transport groups are capable of maintaining consensus even when the nest direction is blocked.

How were groups able to maintain consensus, especially while changing direction? In some cases, direction changes that led to escape seemed to be initiated by new ants joining the transport effort (Table 3). These new ants may have more information about the shape of the obstacle, and will have arrived using a successful path. If the group rapidly conforms to a new individual, as Gelblum *et al.* (2015) reported, they are likely to succeed. Indeed, we found that in 32% of cul-de-sac trials, the initiation of the turn leading to escape coincided with a new ant joining the group. Yet in the remaining trials no new ant was present, and in 58% of all cases we could see no clear cause for the change. For the sharpest turns, which were also common, we again observed that while a substantial portion of turns coincided with new ants joining the transport group (32%), in almost half of cases (45%) we could detect no event precipitating the change. Yet these changes occurred rapidly without stalling. We concur with the conclusion of Gelblum *et al.* (2015) that these groups rapidly conform to newly imposed directions, and we add that in the case of obstacle navigation, these new directions do not need to come from new group members. Existing group members may also impose new directions. They may do so randomly, or because of new information they have received, perhaps from unladen ants not interacting with the object. A highly conforming group may rapidly change direction based on cues from a single individual, whether or not that individual has recently joined the group.

Groups changed direction frequently, and many cases had no clear external cue for doing so. In some cases these direction changes appeared counter-productive, as they occurred just before the group would have reached the end of the wall (Figs. 2A,B). Could occasional spontaneous direction changes be beneficial? Assuming groups do not know the shapes of obstacles they are navigating, direction changes allow for flexibility and could prevent extraordinarily long navigation times. Consider an obstacle shaped like a very long wall. A group may happen to encounter this obstacle close to the left end, but initially turn right to navigate around it. If they never change direction they will have to traverse nearly the entire length of the wall to find the end. On the other hand, if they spontaneously change direction they will find the end relatively quickly. Thus, these spontaneous direction changes allow groups to abandon tactics that appear to be unsuccessful and try new, potentially fruitful, directions.

In addition to spontaneous direction changes, we were able to observe other strategy elements used by *P. longicornis.* Both obstacles require symmetry breaking: after encountering the obstacle groups must choose a travel direction among equal options. In every trial, groups quickly broke symmetry. Groups also followed the perimeter of the obstacle – with few exceptions groups navigating the wall remained close to the obstacle (Fig. S2). This may simply be a result of groups being unlikely to move counter to their preferred direction. Likewise, groups in the cul-de-sac initially stayed close to the back wall, but typically did not continue following the perimeter to travel “backwards” along the sides of the obstacle. Yet the longer a group was in the cul-de-sac, the more time they spent moving backwards, away from the nest. Even then, spontaneous direction changes were always present; direction changes were initially constrained mainly to movement along the back wall, while later on, direction changes were less constrained resulting in a more complete exploration of the space.

These observations give us a partial picture of the group strategy, which initially includes symmetry breaking, moving toward the nest when possible – resulting in perimeter following, and spontaneous direction changes. If the group remains unsuccessful, their behavior changes to include *less* perimeter following and incorporating a new element, which is moving “backwards.” By investigating how these elements are implemented in the groups, future studies could elucidate more about the mechanisms of collective problem solving.

We also investigated the effect of group size on transport and navigation. We found that larger groups moved faster than smaller groups, which agrees with analyses presented by Gelblum *et al.* (2015). Surprisingly, while larger groups had faster speeds, the increase in speed did not result in faster obstacle navigation. In fact, for the cul-de-sac, the mean speed of a group was positively correlated with sinuosity, indicating that faster groups are less efficient at navigating this obstacle. Specific colonies tended to be either relatively slow with low sinuosities or relatively fast with high sinuosities (Fig. 6B). While we only studied a total of four colonies, it is interesting that our two “slower” colonies were those at the Biosphere 2 facility, and our two “faster” colonies were those at Arizona State University. It would be interesting to see whether these differences result simply from colony size or activity level, or indicate more substantial genetic or environmental differences.

While most of our analysis focused on the wall and the cul-de-sac, we also found some unexpected behavior for groups in the trap. The shape of the cul-de-sac and trap only differ by whether or not there is an exit. We expected groups in the trap to behave similarly to groups in the cul-de-sac for at least as long as it took to navigate the cul-de-sac, after this time it is reasonable to imagine that groups may lose energy or stop trying. Instead, we found that groups have dramatically different behavior in the trap, and the difference is apparent right away. Groups slowed down quickly, and we observed individuals spending less and less time grasping the object being carried. This suggests that these ants are capable of detecting that they are trapped, or at least detecting a difference in their situation compared to the cul-de-sac.

While no strategy will successfully navigate the trap, *P. longicornis* workers have a strategy that is robust, in that they do not attempt to solve it endlessly. In natural scenarios, abandoning the navigation effort allows individuals to transition to other behaviors, such as searching more effectively for an exit, or escaping to return to the nest without the object they were carrying. How do ant groups distinguish between the trap and the cul-de-sac? While the trap keeps the group stuck inside, it also prevents other ants from entering the obstacle and interacting with the transport group. Perhaps these additional ants are important to the navigation process. Indeed, Czaczkes *et al.* (2013) suggested that escort ants play an important role in cooperative transport in *P. longicornis,* especially for transport of live prey. These escorts may also be important with non-live prey when navigating obstacles.

Overall, our results demonstrate that *P. longicornis* employ a problem solving strategy that adapts to the complexity of the problem. Future work should aim to understand how this strategy is implemented. How do groups maintain consensus? Perhaps a quorum is required for decisions, as in nest site selection in honey bees and *Temnothorax* (Pratt, 2005; Seeley, 2010), or perhaps informed individuals play an outsized role in decisions (Couzin et al., 2005; Gelblum et al., 2015). How is information integrated and transferred within the group? Information may be transferred through the object itself (Kube and Bonabeau, 2000; McCreery and Breed, 2014), or maybe escorts aid information transfer. This study focused on *P. longicornis;* other ant species that are effective at cooperative transport, such as *Novomessor cockerelli* and *Oecophylla longinoda* may have evolved completely different strategies. For example, groups may use visual detection to avoid obstacles or pheromones to lay trails around obstacles. Ants live in complex environments, where debris may create unexpected obstacles, or a large load may get stuck on harsh terrain; being able to find other routes when the nest direction is blocked is crucial for navigation and is likely to be observed in other species. Understanding the range of strategies employed by ant species would lead to new insights into the information processing and collective problem-solving strategies of ant colonies.

Our study also has general implications for collective problem solving in other animals. Many animals navigate in groups, from schools of fish to large migrations, and these groups must form consensus to remain cohesive. We show that a group reliant on consensus can collectively make a series of stochastic decisions in order to navigate a complex, unknown environment. In addition, obstacle navigation strategies in *P. longicornis* may provide inspiration for the design of new stochastic strategies for robotics (Bonabeau and Théraulaz, 2000). Ant collective behavior has been a rich source of concepts for research in distributed algorithms, including for cooperative transport by groups of robots (Berman et al., 2011; Rubenstein et al., 2013), and insights from highly effective social groups like *P. longicornis* could provide new ideas for designing robot teams that robustly tackle complex environments. Indeed, we show that *P. longicornis* groups are able to collectively navigate complex environments by using a cohesive, flexible, and robust navigation strategy.

## Acknowledgements

We thank James McLurkin for providing some early suggestions about the study framework. Stephen Pratt at Arizona State University and John Adams at Biosphere 2 provided assistance and use of their facilities during fieldwork. We used Justin Werfel's code for data extraction, and Kathleen Kurtenbach, Robert Mason Woodside, and Jenna Bilek helped extract data. We thank Maxwell Joseph for his help implementing the Bayesian analysis in Stan, and for making helpful suggestions about analyses. We thank QDT, modeling group, and the writing coop for providing suggestions and comments on the analyses and writing. The University of Colorado Arts and Sciences Graduate School and the Ecology and Evolutionary Biology Department provided funding.

### Author Contributions

H.F.M., R.N., and M.D.B. designed the experiments. H.F.M. and Z.D. conducted the fieldwork and contributed to extracting data from videos. H.F.M. and R.N. interpreted data. H.F.M. conducted statistical analyses and wrote the article, with editing contributed from all authors.

